# Transcriptome analysis of the NR1H3 mouse model of multiple sclerosis reveals a pro-inflammatory phenotype with dysregulation of lipid metabolism and immune response genes

**DOI:** 10.1101/2021.03.17.435678

**Authors:** Carles Vilariño-Güell, Mary Encarnacion, Cecily Q Bernales, Emily Kamma, Pierre Becquart, Jacqueline A Quandt

## Abstract

**Background:** The development of effective treatments for multiple sclerosis (MS), and in particular its progressive forms, is hampered by the lack of etiologically relevant cellular and animal models of human disease. Models that recapitulate the biological and pathological processes leading to the onset and progression of MS in patients are likely to afford better translational efficacy. Following the discovery of the NR1H3 p.Arg415Gln pathogenic mutation for progressive MS in two Canadian families, we developed a knock-in mouse model harboring a homologous mutation in the endogenous gene to provide a more physiologically relevant model of human MS.

**Methods:** Gene expression was evaluated in constitutive heterozygote (which recapitulates the human disease genotype) and homozygote Nr1h3 p.Arg413Gln knock-in mice on a C57BL/6 background, and compared to wild-type littermates. AmpliSeq Transcriptome Mouse Gene Expression kits analyzed on an Ion Proton sequencer were used to generate the gene expression profiles of spleen, liver, brain and spinal cord tissue from three-month-old male and female mice. Differential expression between genotypes was assessed with DESeq2, and Gene Ontologies pathways enrichment analysis performed with DAVID v6.8. Benjamini-Hochberg false discovery rate (FDR) correction for multiple testing was applied.

**Results:** Transcriptome analysis of spleen tissue from Nr1h3 p.Arg413Gln mice revealed 23 significantly dysregulated genes (FDR<0.05) with greater than a two-fold change in expression. These include CD5 antigen-like (*Cd5l*), complement component 6 (*C6*), procollagen C-endopeptidase enhancer 2 (*Pcolce2*), interleukin 22 receptor, alpha 2 (*Il22ra2*), and T cell immunoglobulin and mucin domain containing 4 (*Timd4*). Gene Ontology enrichment analysis support upregulation of cell cycle pathways and downregulation of immune system response in splenic cells. The liver transcriptome identified 27 significantly dysregulated genes with greater than a two-fold change in expression compared to wild-type littermates. *Cd5l*, *Timd4*, C-C motif chemokine receptor 3 (*Ccr3*), ADAM metallopeptidase domain 11 (*Adam11*) and macrophage expressed 1 (*Mpeg1*) were amongst those most significantly dysregulated. Enrichment analysis supported altered immune function with upregulation of sterol and steroid metabolic processes and downregulation of fatty acid biosynthesis and inflammatory and immune system responses. Although brain and spinal cord transcriptome profiles identified several genes significantly dysregulated in *Nr1h3* mice compared to wild-type littermates (FDR<0.05), none presented greater than two-fold changes in gene expression.

**Discussion:** The analysis of the Nr1h3 p.Arg413Gln mouse model of MS suggests that the predominance of a pro-inflammatory over a healing or reparative phenotype, combined with deficiencies in myelination and remyelination, are the biological mechanisms implicated in the onset of MS and the development of a more severe progressive disease course observed in patients with *NR1H3* mutations. Association of *NR1H3* common variants with MS risk indicates that the disruption of these biological and immunological processes is not only informative for familial forms of disease but MS patients at large. Differences in transcriptome profiles underline the value of this model for the development and validation of novel therapeutic strategies and ultimately treatments with the potential to delay or even halt the onset of progressive MS and to ameliorate the severity of clinical symptoms.

## INTRODUCTION

Multiple sclerosis (MS) is a common inflammatory disease characterized by myelin loss, varying degrees of axonal pathology, and progressive neurological dysfunction. An increasing number of disease-modifying treatments for MS have been approved since the 1990s as a result of a better understanding of the biological mechanisms implicated in disease. However, available treatments only modestly impact new disease activity, and their usage can present several serious or even life-threatening adverse effects (1). A substantial contribution to our understanding of the biological mechanisms in MS has been provided by the commonly used experimental autoimmune encephalomyelitis (EAE) animal model. Although EAE has provided insight into the mechanism of autoimmunity, neuroinflammation and cytokine biology, it fails to provide insight into disease progression (2). While toxic models of demyelination and remyelination have been used to try to identify strategies to correct the progressive and neurodegenerative phase of MS, they lack the extensive immune activity seen in MS patients (2). It is therefore imperative to develop cellular and animal models of MS which recapitulate human disease etiology to better understand the biological processes triggering the onset of disease and progression in people living with MS, and to provide reliable and accurate translational efficacy for the identification of more effective treatments.

To fulfill this need, we have generated a knock-in mouse model harboring a substitution in its endogenous nuclear receptor subfamily 1 group H member 3 (Nr1h3 p.Arg413Gln) gene that is homologous to the pathogenic mutation (p.Arg415Gln) identified in two multi-incident MS families presenting primary progressive (PP)MS or a rapidly progressive disease course (3). *NR1H3* encodes liver X receptor alpha (LXRA), a nuclear receptor that controls the transcriptional regulation of genes involved in lipid homeostasis, inflammation, and innate immunity (4). The mutation identified in MS patients was shown to impair the dimerization of LXRA with retinoid X receptors, altering the transcriptional regulation of its cis targets. LXRA has also been shown to regulate gene expression through a process known as transrepression (5), a mechanism which does not appear to be disrupted by the p.Arg415Gln mutation (3) thus making our model distinct to previously described LXRA null mice (6).

It is important to note that significant associations between common variants in *NR1H3* and patients presenting PPMS have been described, and were replicated in unstratified large-scale association studies of MS (3,7). In addition, a mutation in nuclear receptor coactivator 3 (*NCOA3*), a critical element of the NR1H3 nuclear receptor activation complex, was found to co-segregate with disease in multi-incident MS families, indicating that the nuclear receptor pathways may be an important contributor to MS development (8). These studies suggest that the characterization of our mouse model could provide a better understanding of the pathways leading to the onset of MS in families with rare *NR1H3* mutations, but also has the potential to elucidate the pathomechanisms of disease in patients with other rare genetic mutations as well as patients without a family history of MS. Given the predominantly progressive disease course observed in *NR1H3* mutation carriers, and association of common genetic variants with PPMS patients, characterization of this model could shed light on the biological mechanisms implicated in the development of the highly debilitating and largely treatment-resistant progressive phase of the disease.

## METHODS

### Animals

Conditional Nr1h3^R413Q/+^ C57BL/6 mice were generated by GenOway Inc (Lyon, France) and crossed with C57BL/6 Tie2-Cre mice (9) to create constitutive Nr1h3 p.Arg413Gln heterozygote and homozygote knock-in mutant mice (**Fig. 1**). PCR product size discriminates between conditional and constitutive mice, as well as between genotypes which were confirmed by Sanger sequencing. No significant differences in *Nr1h3* expression were observed between genotypes in any of the tissues examined (**Fig S1**). Gene expression profiles were generated from constitutive homozygote (Nr1h3^R413Q/R413Q^) and heterozygote (Nr1h3^R413Q/+^) mice, and compared to wild-type littermates (Nr1h3^+/+^).

**Fig 1.**
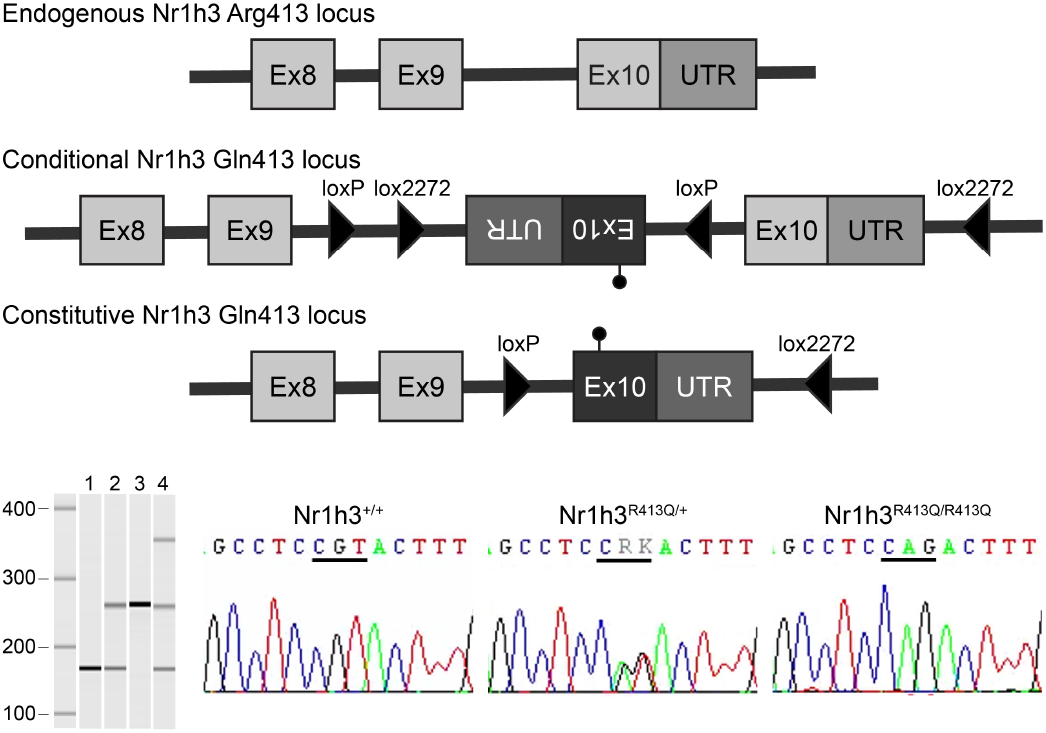
Schematic of the *Nr1h3* locus in wild-type, conditional and constitutive *Nr1h3* mice with the p.Arg413Gln mutation indicated with a flag; PCR size discrimination of wild-type mice (1), and constitutive heterozygote (2), constitutive homozygote (3) and conditional heterozygote (4) *Nr1h3* knock-in mice; and representative electropherograms for each constitutive genotype.

### Transcriptome analysis

Total RNA was extracted from spleen, liver, brain and spinal cord flash frozen tissues using standard protocols.

Tissue samples were studied from nine to twelve three-month-old mice of each sex and genotype for spleen, liver and brain, and three mice of each sex and genotype for spinal cord. Whole transcriptome sequencing was performed using 100ng of total RNA with Ion AmpliSeq Transcriptome Mouse Gene Expression kits on an Ion Proton sequencer (Life Technologies, Carlsbad, CA), with an average of 11.7 (SD ± 2.9) million reads per sample. The Ion Torrent Server (v5.8) was used to map reads to the GRCm38 (mm10) reference genome, and the AmpliseqRNA plugin to generate absolute gene expression reads. Differential expression between genotypes was quantified using DESeq2 (v1.22.0) lfcShrink on genes with a minimum of 10 reads across samples (10). Differences in gene expression between wild-type and heterozygote or homozygote genotypes were evaluated for each tissue separately using a Wald test. Male and female mice were analyzed together unless principal component analysis (PCA) indicated differential expression profiles between sexes. Enrichment analysis of Gene Ontologies pathways for differentially expressed genes was calculated using the Database for Annotation, Visualization and Integrated Discovery (DAVID v6.8) (11). Benjamini-Hochberg false discovery rate (FDR) correction for multiple testing was used for mRNA expression profiling and pathway analysis. Differential gene expression for five significantly dysregulated genes in each analysis was reassessed and confirmed in six randomly selected samples of matching sex and genotype using SYBR Green on an Applied Biosystems 9700 Real-Time PCR System and normalized to the geometric mean expression of three housekeeping genes (*Actb*, *Gapdh* and *Rpl19*) (data not shown).

## RESULTS

Gene expression profiles from three-month-old mice of each sex and genotype were generated for the identification of transcriptional differences between heterozygote (Nr1h3^R413Q/+^) or homozygote (Nr1h3^R413Q/R413Q^) constitutive knock-in mice and wild-type littermates (Nr1h3^+/+^). The analysis of heterozygote mice recapitulates the genotype observed in MS families, whereas homozygote mice were considered likely to cause a more overt phenotype emphasizing the genes and pathways most disrupted by the *Nr1h3* mutation.

PCA identified distinct patterns of gene expression between sexes in liver, brain and spinal cord tissue (**Fig. S2**). Thus gene expression in spleen tissue was determined for both sexes combined, whereas transcriptome analyses for all other tissues were performed for each sex separately. Differentially expressed genes, defined as those presenting statistically significant differences (FDR <0.05) with a minimum of two-fold change in gene expression (log_2_ fold (L_2_F) >|1|) compared to wild-type littermates, were observed in spleen and liver tissues from *Nr1h3* knock-in mice (**Fig. 2**). Although statistically significant differences between genotypes were observed in brain and spinal cord tissues from *Nr1h3* mice, those genes presented less than a two-fold expression change compared to wild-type mice (**Fig. S3 & S4**).

**Fig 2.**
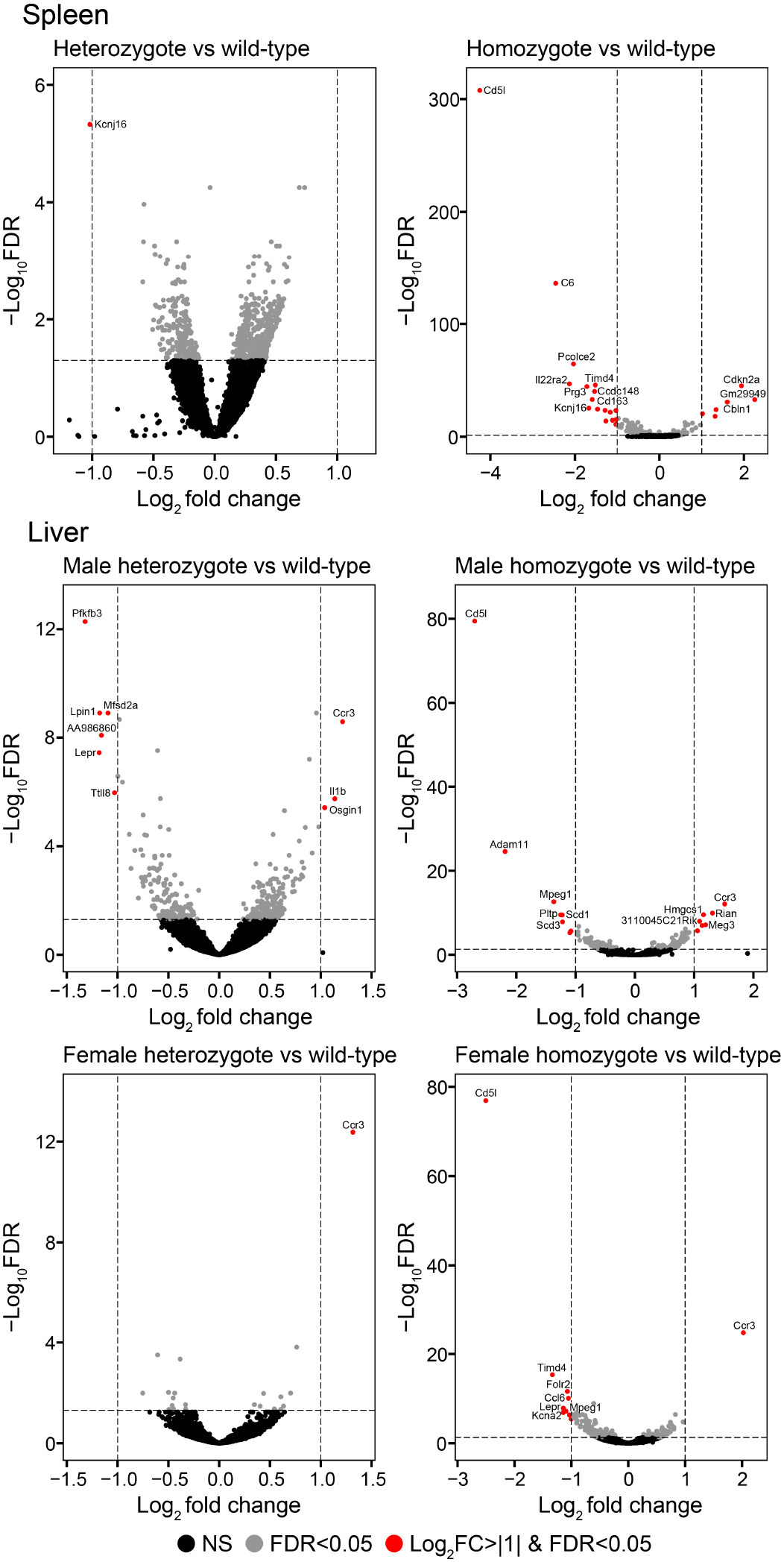
Volcano plots for spleen and liver tissue from heterozygote and homozygote Nr1h3 p.Arg413Gln mice.

### Spleen transcriptome profile

A total of 23 genes were found to be significantly dysregulated in homozygote Nr1h3 p.Arg413Gln knock-in mice compared to wild-type littermates (**Fig. 2**). The genes most significantly dysregulated are CD5 antigen-like (*Cd5l*, L_2_F=−4.25), complement component 6 (*C6*, L_2_F=−2.45), procollagen C-endopeptidase enhancer 2 (*Pcolce2*, L_2_F=−2.03), interleukin 22 receptor, alpha 2 (*Il22ra2*, L_2_F=−2.12), and T cell immunoglobulin and mucin domain containing 4 (*Timd4*, L_2_F=−1.52). The majority of genes found to be differentially expressed in homozygote mice are also significantly dysregulated in heterozygote *Nr1h3* mice (FDR<0.05). However, with the exception of potassium inwardly rectifying channel subfamily J member 6 (*Kcnj6*, L_2_F=−1.02), they all presented less than a 2-fold change in expression (L_2_F<|1|) in heterozygote mice relative to wild-type littermates (**Fig. 3**). This data suggests that gene expression in the spleen of heterozygote p.Arg413Gln mice largely represent an intermediate phenotype between wild-type and homozygote carriers.

**Fig 3.**
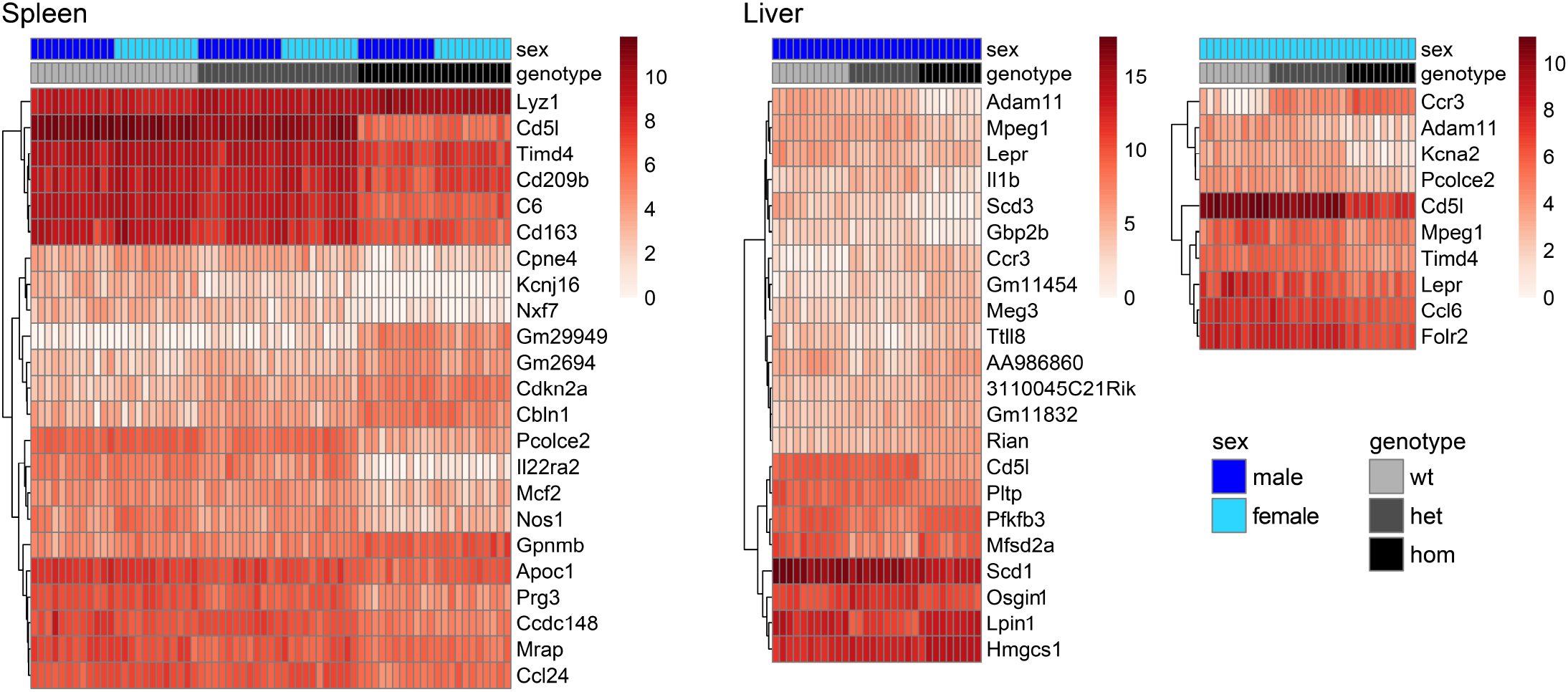
Hierarchical clustering of normalized transform gene expression (log_2_(n+1)) for significantly differentially expressed genes (FDR<0.05) with a minimum of two-fold change (L_2_F>|1|) in spleen or liver tissue of *Nr1h3* mice compared to wild-type littermates.

To identify the biological processes associated with the transcription profiles observed in *Nr1h3* mice we cross-referenced all significant genes (FDR<0.05) with the Gene Ontology database for biological processes (**Fig. 4**). Upregulated genes in *Nr1h3* knock-in mice were found to be enriched for processes involving DNA replication and cell cycle and division; downregulated genes were enriched for biological pathways involving regulation of transcription and immune system response.

**Fig 4.**
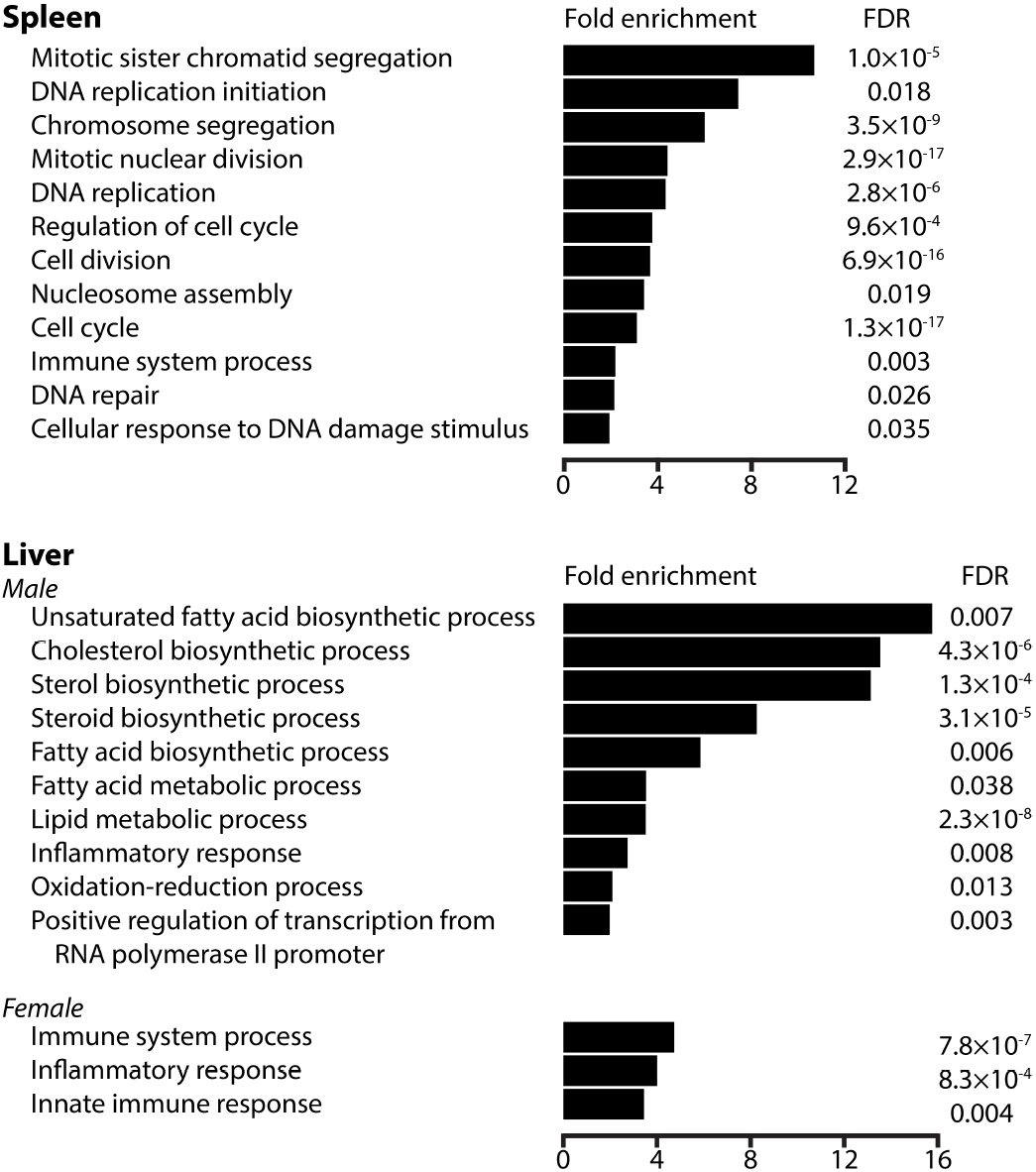
Gene Ontology (GO) enrichment analysis for differentially expressed genes (FDR<0.05) in spleen or liver tissue.

### Liver transcriptome profile

A total of 27 genes were found to be differentially expressed in liver tissue from *Nr1h3* mutant mice compared to wild-type littermates (**Fig. 2**). Fourteen genes were found to be dysregulated in male and female homozygote knock-in mice, albeit not always with greater than a 2-fold difference in expression for both sexes. The remaining 13 genes were found to be differentially expressed exclusively in male mice, six in homozygote and seven in heterozygote animals. Genes found to be most significantly dysregulated in male mice are *Cd5l* (L_2_F=−2.70), ADAM metallopeptidase domain 11 (*Adam11*, L_2_F=−2.19), macrophage expressed 1 (*Mpeg1*, L_2_F=−1.37), C-C motif chemokine receptor 3 (*Ccr3*, L_2_F=1.52), and 6-phosphofructo-2-kinase/fructose-2,6-biphosphatase 3 (*Pfkfb3*, L_2_F=−1.32). In female mice, the most significantly differentially expressed genes are *Cd5l* (L_2_F=−2.50), *Ccr3* (L_2_F=2.03), *Timd4* (L_2_F=−1.33), folate receptor beta (*Folr2*, L_2_F=−1.07), and chemokine (C-C motif) ligand 6 (*Ccl6*, L_2_F=−1.05).

Although the majority of genes found to be differentially expressed in homozygote mice were not significantly dysregulated in heterozygote mice, they largely present an intermediate expression level between homozygote and wild-type animals, thus suggesting a more moderate phenotype for heterozygote carriers (**Fig. 3**). In contrast, the seven genes found to be dysregulated exclusively in heterozygote mice present a gene expression profile in homozygote mice that is largely comparable to wild-type littermates. This suggests the Nr1h3 p.Arg413Gln substitution causes a heterozygote specific phenotype in the liver that is particularly apparent in male mice.

Gene Ontology biological pathway analysis identified an enrichment of lipid metabolic processes in male mice (**Fig. 4**). Upregulated genes were enriched in cholesterol, sterol and steroid biosynthesis and metabolic processes, whereas downregulated genes were enriched in fatty acid metabolic and biosynthetic processes. Downregulated genes were further enriched in inflammatory response and immune system processes in male and female mice.

### Brain and spinal cord transcriptome profile

Transcriptome analysis of whole brain or spinal cord tissues from three-month-old *Nr1h3* mice failed to identify genes with greater than 2-fold change in expression relative to wild-type mice (**Fig. S3 & S4**). However statistically significant differences with a minimum of 1.3-fold change in gene expression were observed in brain for serum/glucocorticoid regulated kinase 1 (*Sgk1*, L_2_F=−0.85), BTG anti-proliferation factor 2 (*Btg2*, L_2_F=0.58), iodothyronine deiodinase 2 (*Dio2*, L_2_F=−0.40), immediate early response 2 (*Ier2*, L_2_F=0.47), and neuronal PAS domain protein 4 (*Npas4*, L_2_F=0.40) in heterozygote male mice, and carbonic anhydrase 5b, mitochondrial (*Car5b*, L_2_F=0.44) in homozygote females. Similarly, significant expression changes with greater than 1.3-fold difference in expression compared to wild-type littermates was observed for *Sgk1* (L_2_F=−0.75) and solute carrier family 30 member 3 (*Slc30a3*, L_2_F=0.72) in spinal cord tissue from heterozygote and homozygote male mice respectively.

## DISCUSSION

The availability of appropriate mouse models of MS, capable of recapitulating the biological pathways that influence disease development in patients is essential to gain a better understanding of the pathomechanisms of MS. In this study we analyzed spleen, liver, brain and spinal cord transcriptome profiles for the first animal model of MS based on human genetic etiology (3).

Compromised immune function is depicted by reductions in *Cd5l*, *Timd4*, *C6*, *Il22ra2*, *Pfkfb3* and upregulation of *Ccr3* and interleukin 1 beta (*Il1b*) amongst others. Similarly, dysregulation of lipid biosynthesis and metabolism is underlined by differential expression of *Pcolce2*, apolipoprotein C1 (*Apoc1*), stearoyl-coenzyme A desaturase 1 (*Scd1*), phospholipid transfer protein (*Pltp*), and lipin 1 (*Lpin1*) amongst others.

*Cd5l* and *Timd4* were markedly downregulated in both spleen and liver tissues from *Nr1h3* mice (**Fig. 3**). CD5L is a secreted glycoprotein that modulates several key inflammatory responses. CD5L polarizes activated macrophages to express markers and factors associated with immunomodulation, healing and repair (12); this is in contrast to a more proinflammatory profile that is typically driven by pathogens. In both EAE and MS, macrophages typically change this inflammatory profile over the course of disease onset and recovery (13,14). CD5L also acts as a functional switch to regulate T-helper (Th)17 cell pathogenicity; its loss has been shown to convert otherwise non-pathogenic Th17 cells that also produce the immunomodulatory interleukin-10 cytokine into pathogenic cells in models of autoimmunity (15). TIMD4 is a phosphatidylserine receptor that enhances the engulfment of apoptotic cells, a primary role of macrophages in response to tissue injury. Activation of TIMD4 promotes Th1 cell apoptosis and Th2 cell proliferation (16,17), and has a prominent role in autoimmune diseases including rheumatoid arthritis and systemic lupus erythematosus (18).

Decreased expression of the complement component C6 was observed in spleen tissue from heterozygote and homozygote *Nr1h3* mice. C6 is a constituent of the membrane attack complex (MAC), a key contributor to innate and adaptive immune responses implicated in immune-mediated neurodegeneration (19). Deficiencies in MAC formation in C6 knock-out mice result in delayed clearance of myelin debris in post-traumatic peripheral nerve injury, and unexpectedly, improved axonal regeneration and functional recovery (20,21). While these scenearios might link C6 depletion to improved outcomes in autoimmunity, complement proteins also have non-inflammatory roles in the developing brain and synapse loss in neurodegenerative disease, contributing to progenitor cell proliferation, neuronal migration, and synaptic pruning (22).

*Il22ra2* is one of the genes containing genetic variants strongly associated with increased MS risk (7). *Il22ra2* expression was significantly reduced in spleen from *Nr1h3* mice, but has been found to be overexpressed in monocytes and dendritic cells from MS patients (23). Similarly, IL22RA2 deficiency results in less demyelination and milder clinical disease following the induction of EAE, but exacerbated skin inflammation after the induction of imiquimod-induced skin disease (23,24), suggesting pleiotropic effects.

The chemokine receptor CCR3 is expressed on basophils, mast cells, T cells, dendritic cells, brain microglial cells, epithelial cells and endothelial cells (25). While typically linked to immune cell recruitment, increasing evidence support CCR3 roles in repair and vascular remodeling (26), warranting histological examination.

Other genes affecting immune profiles identified in this study include reductions in markers of reparative macrophage polarization CD163 antigen, CD209b antigen and FOLR2 (27,28); the CCR3 ligand eotaxin-2 (CCL24) (29); the macrophage expressed perforin-2 (*Mpeg1*), a bacterial cell wall pore-forming protein with roles in innate immunity (30); and increased expression of the negative regulator of macrophage inflammation GPNMB (31).

Heterozygote-specific differences were observed in *Nr1h3* male mice, those include increased *Il1b* and decreased *Pfkfb3* expression in liver (**Fig. 2**). IL1B is a potent pro-inflammatory cytokine with broad immune functions, including B- and T-cell activation and Th17 differentiation. Elevated IL1B expression has been described for several autoimmune disorders as well as PPMS patients (32,33), the predominant disease course observed in NR1H3 p.Arg415Gln mutation carriers. *Pfkfb3* encodes a critical glycolysis-regulatory enzyme required for cell cycle progression and prevention of apoptosis, and protects against diet-induced intestine inflammation (34). Inhibition of Pfkfb3 in cardiomyocytes suppresses the secretion of LPS-mediated pro-inflammatory cytokines and diminishes the nuclear translocation and phosphorylation of tumor necrosis factor-alpha (35).

Dysregulation of lipid biosynthesis and metabolic pathways was observed predominantly but not exclusively in male mice (**Fig. 4**). Obesity is a well known risk factor for MS, and is often associated with chronic inflammation (36,37). In *Nr1h3* mutant mice, we identified dysregulation of fatty acid, cholesterol, sterol and steroid biosynthesis and metabolism; biological processes known to be important for myelination and remyelination processes. The formation of myelin sheaths, which are composed of 70-85% lipids and with high cholesterol content, require activation of LXR pathways, high levels of fatty acid and lipid synthesis, and uptake of extracellular fatty acids (38,39).

In *Nr1h3* mutant mice we observed reduced expression of *Pcolce2* in both liver and spleen (**Fig. 3**). PCOLCE2 has a pivotal role in high-density lipoprotein (HDL) metabolism, reverse cholesterol transport, and atherosclerosis (40). Null PCOLCE2 mice revealed impaired ability to mediate in-vitro cholesterol efflux via ABCA1, resulting in elevated concentrations of enlarged HDL particles (41).

APOC1 regulates the activity of enzymes and receptors involved in lipid and cholesterol metabolism, modulating a variety of biological processes affecting inflammation, immunity and cognition amongst others. In this study, we observed decreased *Apoc1* expression in spleen from *Nr1h3* mice. *Apoc1* is also expressed in astrocytes and endothelial cells from the human hippocampus, as well as during monocyte differentiation into macrophages and the formation of foam cells (42). *Apoc1* knock-out mice were found to have impaired hippocampal-dependent memory function and increased concentrations of pro-inflammatory markers (43).

SCD1 is a central and rate-limiting lipogenic enzyme in the synthesis of monounsaturated fatty acids from saturated long-chain fatty acids. SCD1 has essential roles in the regulation of lipid synthesis and oxidation, hormonal signaling, and inflammation (44). SCD1 knock-out animals present decreased expression of liver lipogenic genes and increased plasma levels of insulin and leptin, resulting in resistance to diet-induced obesity and protection from liver injury (45,46). Phospholipid transfer protein (PLTP) is a crucial protein in reverse cholesterol transport that facilitates the transfer of phospholipids from triglyceride-rich lipoproteins into HDL. Similarly to SCD1 deficient mice, PLTP knock-out animals show reduced high fat diet-induced obesity. In addition, they presented decreased levels of sphingomyelin and free cholesterol (47).

Liver tissue from heterozygote male mice revealed the dysregulation of several lipid metabolic genes, most notably lipin 1 (*Lpin1*), that were not affected in *Nr1h3* female mice or homozygote male mice (**Fig. 2**). LPIN1 is a phosphatidic acid phosphatase enzyme that controls the metabolism of fatty acids. *Lpin1* knock-out mice develop lipoatrophy and progressive demyelinating neuropathy affecting peripheral nerves (48), similar to the adult-onset syndromic myasthenia with peripheral neuropathy phenotype observed in two patients with compound heterozygote *Lpin1* mutation (49). In contrast, recessive loss of function mutations in *Lpin1* are more commonly described in patients presenting early childhood myoglobinuria and rhabdomyolysis associated with muscle pain and weakness (50). Other interesting genes with roles in lipid metabolism and reduced expression in *Nr1h3* mice are leptin receptor (*Lepr*) and *Scd3*. Recessive mutations in *Lepr*, which controls hypothalamic regulation of food intake, cause severe early-onset obesity with T-cell deficiency, and is associated with nonalcoholic fatty liver disease and type-2 diabetes (51–53). SCD3, a stearoyl-CoA desaturase like SCD1, catalyzes the synthesis of monounsaturated fatty acids required for the biosynthesis of membrane phospholipids, cholesterol esters and triglycerides (54).

Transcriptome profiles from brain and spinal cord tissues from *Nr1h3* mutant mice only showed minor differences in gene expression compared to wild-type animals. Interestingly *Sgk1*, a serine-threonine proteinase with roles in neuronal excitability and immune regulation (55,56), was downregulated in brain and spinal cord tissues from heterozygote male mice exclusively, albeit with a 1.74 and 1.59-fold decreased expression compared to wild-type littermates. Similarly, the expression of zinc transporter 3 (*Slc30a3*), which modulates neurogenesis, inflammation and demyelination (57,58), was found to be increased (1.15-fold) in spinal cord from *Nr1h3* homozygote male mice (**Fig. S4**).

The lack of major differences in brain and spinal cord expression profiles was slightly unexpected, given that significant associations between common *NR1H3* genetic variants and MS risk were stronger in patients with PPMS than those with relapsing remitting disease (3). Our data suggests that the neurological dysfunction observed in MS patients may be secondary to the deficiencies in immune response and lipid metabolism processes observed in this model. However, analysis in early stages of brain development as well as single cell transcriptome analysis in adult mice will be necessary to determine whether gene expression differences do exist in brain or spinal cord in specific embryonic stages not assessed in our study, or certain cell types or regions that may be masked by tissue heterogeneity. Further assessment of brain and spinal cord tissues from *Nr1h3* mutant mice is warranted given the differential expression, albeit only in spleen and liver, of genes relevant to neuronal function. Those include: *Kcna2* which contributes to the regulation of neuronal action potentials and with gain of function mutations causing epileptic encephalopathy (59); nitric oxide synthase 1 (*Nos1*) which synthesizes nitric oxide, an important immune modulator known to be implicated in neurotoxicity associated with several neurodegenerative diseases (60); and major facilitator superfamily domain containing 2A (*Mfsd2a*) a sodium-dependent lysophosphatidylcholine symporter essential for normal brain growth and cognitive function, and important for blood-brain barrier formation and function (61,62).

Taken together, transcription profiles from *Nr1h3* knock-in mice suggest that the mechanisms of disease for MS patients harboring the p.Arg415Gln mutation are likely the result of exacerbated inflammatory responses harboring a more pro-inflammatory than anti-inflammatory or reparative phenotype, compounded with deficiencies in myelination and remyelination processes, exacerbating neuronal damage as a result of dysregulation of sterol and fatty acid metabolism.

In conclusion, identifying a pathogenic mutation for familial MS in *NR1H3* has led to the generation of the first physiological model of human disease for MS. The Nr1h3 p.Arg413Gln mouse model herein described provides the means for identifying and characterizing the biological pathways responsible for the onset of MS in these families. Importantly, the identification of risk alleles for MS, and particularly for patients presenting PPMS course, ensures that the discoveries from this model will be relevant to a large proportion of MS patients, beyond those with very rare mutations in *NR1H3* or related genes (3,8). Gained insight from this model is expected to provide novel biological targets for therapeutic intervention, particularly for the progressive phases of the disease, and the means for developing and assessing the efficacy of these new treatments.

## ACKNOWLEDGMENTS

This research was undertaken thanks to funding from the Michael Smith Foundation for Health Research (16827), the Vancouver Foundation - The Peggy Griffin Memorial Fund (ADV14-1597), the National Multiple Sclerosis Society (PP-1709-29161), the Multiple Sclerosis Society of Canada (3852) and the VGH and UBC Hospital Foundation Neuroprotection Fund.

