## Supplemental material for "Transcriptome analysis of the NR1H3 mouse model of multiple sclerosis reveals a pro-inflammatory phenotype with dysregulation of lipid metabolism and immune response genes"

**Figure S1.** *Nr1h3* expression (normalized read counts) in spleen, liver, brain and spinal cord tissue from Nr1h3$^{+/+}$ (wt), Nr1h3$^{R413Q/+}$ (het), and Nr1h3$^{R413Q/R413Q}$ (hom) mice.

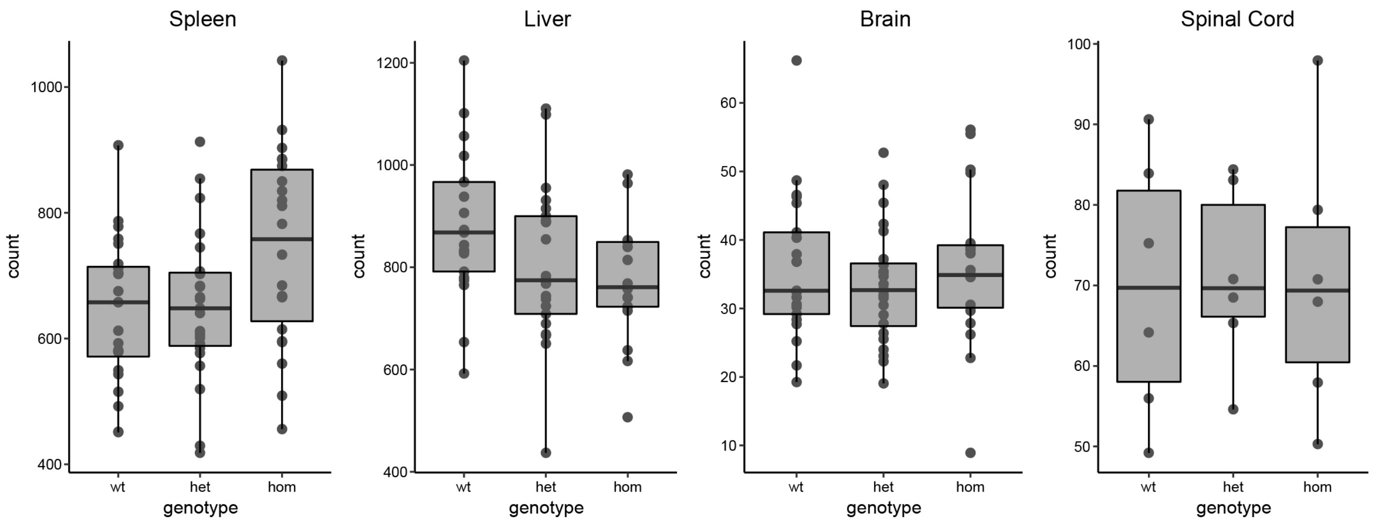

**Figure S2**. Principal component analysis for spleen, liver, brain and spinal cord transcriptome data from three month old mice.

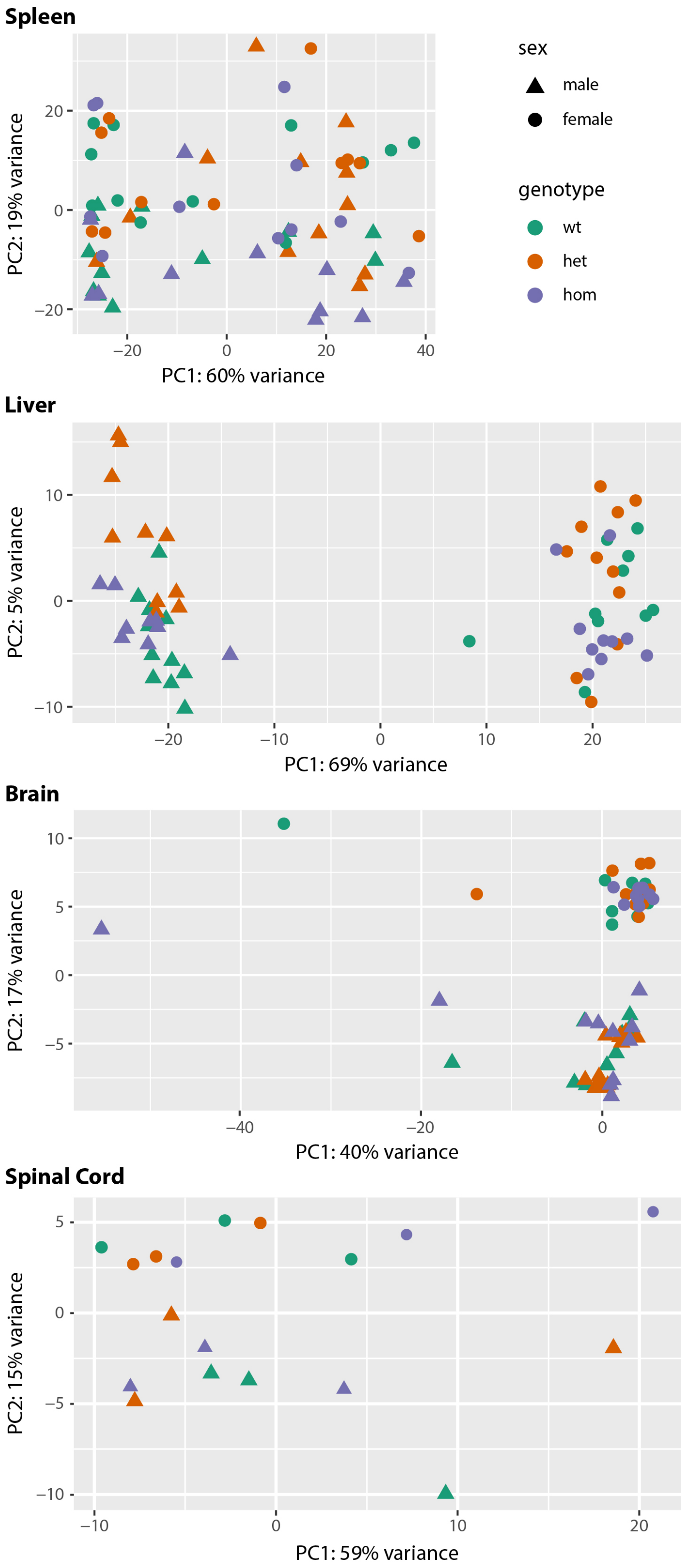

**Figure S3**. Volcano plots for brain tissue from heterozygote and homozygote Nr1h3 p.Arg413Gln mice.

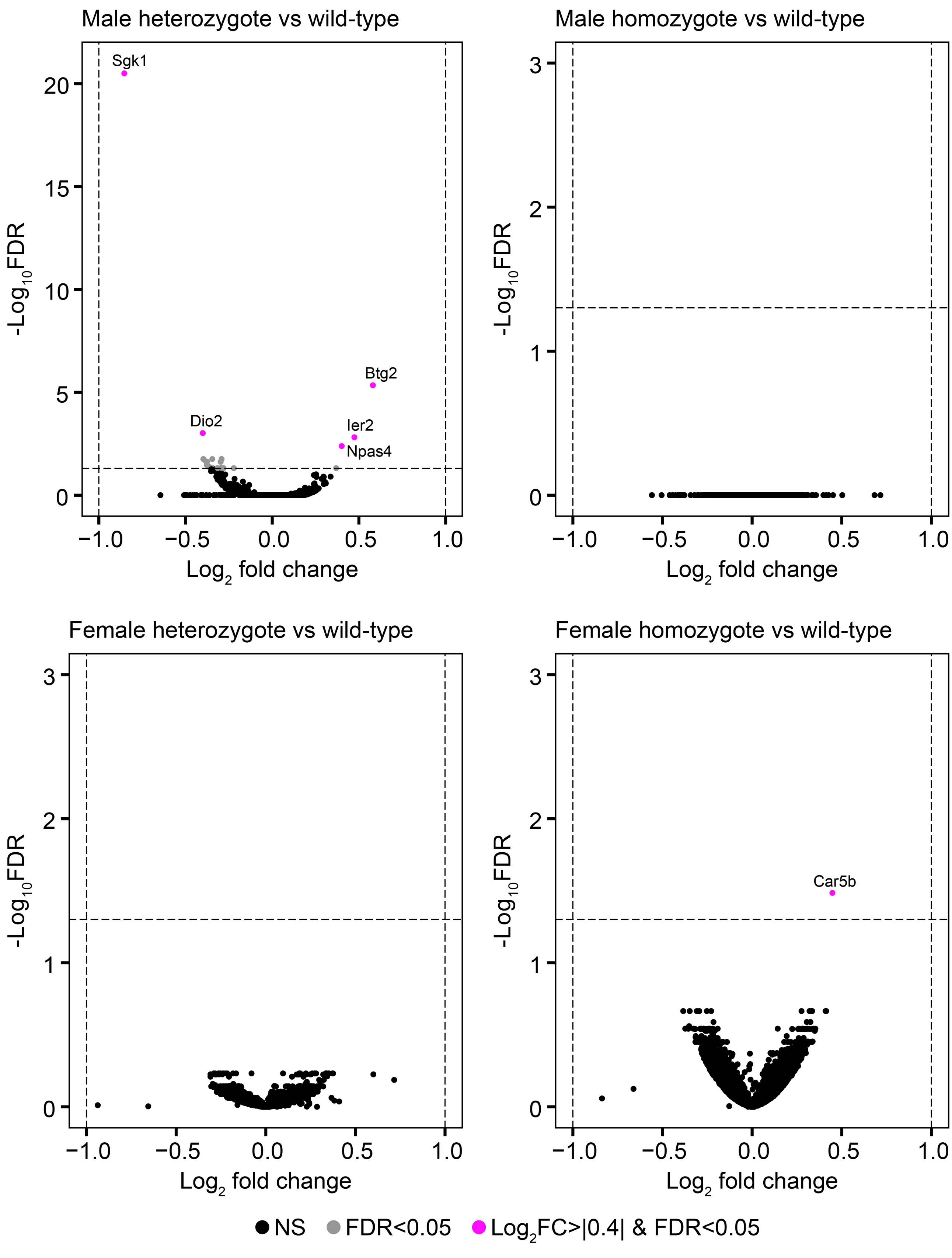

**Figure S4**. Volcano plots for spinal cord from heterozygote and homozygote Nr1h3 p.Arg413Gln mice.

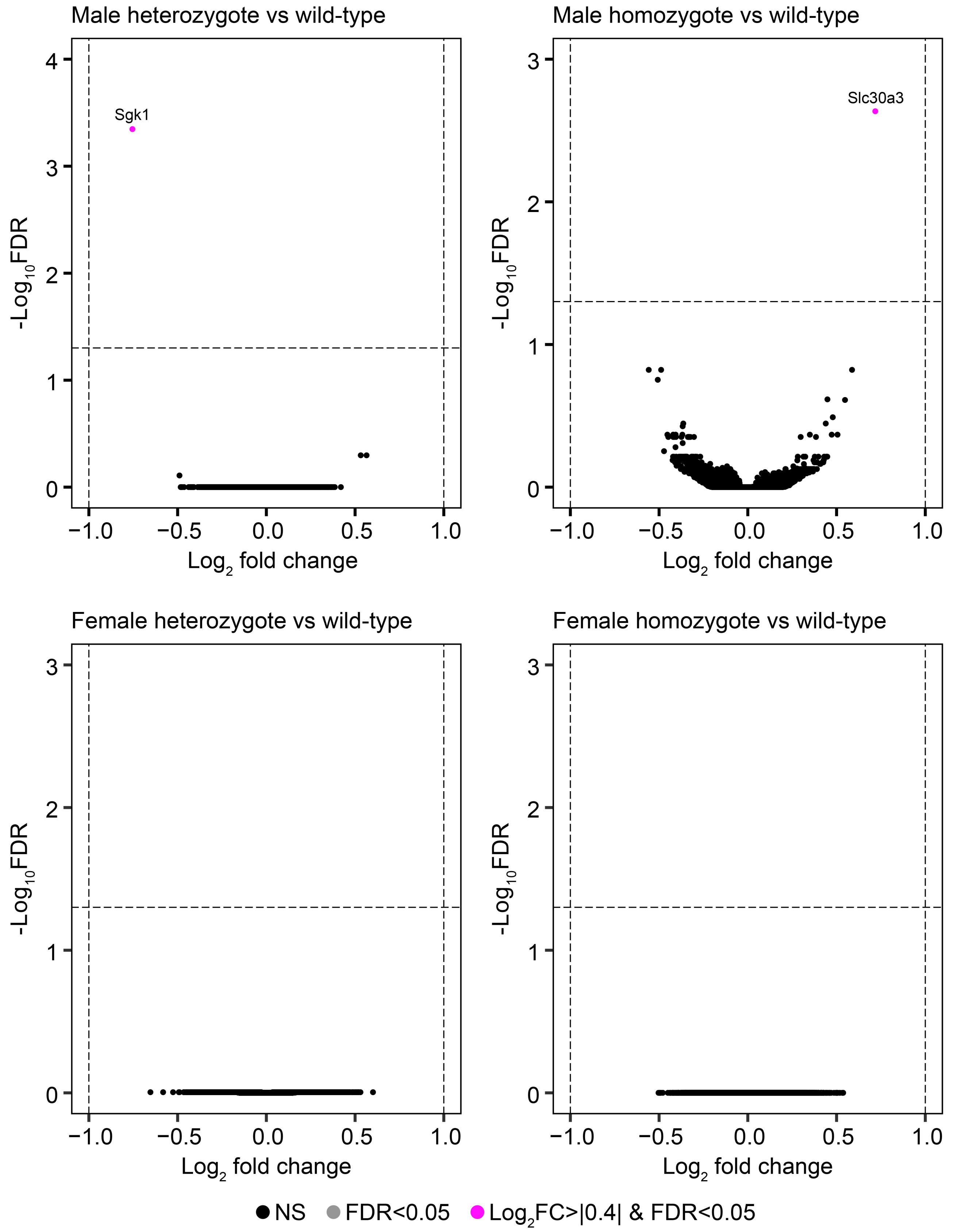
